# Provenance-aware handling of identical dengue genome records alters inferred envelope-protein variation in Bangladesh DENV-3

**DOI:** 10.64898/2026.11.04.764337

**Authors:** Kazi Tanzina Begum, Kazi Abdul Gofur Rafea

## Abstract

The reliability of interpreting dengue virus (DENV) envelope (E) protein variations depends on whether identical public sequence records are kept as representatives or eliminated based on provenance reviews. We evaluated three record-handling strategies using a frozen Bangladesh-associated dataset of 426 E sequences: exact-sequence deduplication, a prespecified provenance-sensitive exclusion method, and an independently curated Nextstrain accession list. Within the 2021 DENV3 subset (96 accessions), 18 records dated August 2021 were identical across 10,707 nucleotides to the older Sri Lankan genome AY099336.1. There were 21 variable E positions within the original 96-accession DENV3 cohort. Whole-cohort deduplication (47 retained) and retaining one representative of the 18-record group (79 retained) preserved all 21 positions. Excluding the group (78 retained) resulted in eight variable positions: 13 of 21 (61.9%) were lost due to differences in state at those positions between the excluded group and retained cohort. Nextstrain reference-specific exclusions selected the same 18 records; applying its full saved list produced 77 accessions and the same eight variable positions. E270 and E383, which had pre-2021 antibody-related evidence, did not meet the sequence-variation criteria for follow-up; however, E223 and E224 remained variable. Antibody-position enrichment was not statistically significant. These results indicate that cohort eligibility decisions may impact dengue E-protein variant prioritization despite representative-retaining deduplication. Genome identity does not indicate specimen identity or an antigenic phenotype.

## Introduction

The use of public viral sequence databases has been instrumental in conducting molecular epidemiology, reconstructing evolutionary relationships among viruses, performing comparative genomics, and monitoring viral activity [1]. However, the utility of these databases relies heavily on the quality and reliability of the data contained within each record, including the origin and interpretability of the data. In some cases, multiple records may contain identical sequences due to either legitimate biological sampling, redundant observations, derivative representations of references, or other types of archival and analytical procedures. Treating these records as separate, individual biological data points can have an effect on how we view both sequence frequency distribution, phylogenetic representation, and estimate of genetic diversity. How we handle the duplicate records will determine what the analytical implications are. If we keep them in our sample size then they will be used to represent themselves; if we remove them from our sample size then we will no longer use them to represent anything. It is impossible to define or manage duplicate records without considering the specific biological question being addressed. Chen et al. identified various types of redundancy in both sequence and annotation data and demonstrated that the impact of this redundancy varies depending on the subsequent analysis performed [2]. Therefore, while exact nucleotide identity is a computational finding, it does not provide sufficient evidence to determine whether two accession records represent the same biological sampling event or whether one should be excluded. Dengue virus consists of four antigenically distinct serotypes. Of particular interest is the variation in the envelope protein (E), as E is the primary surface glycoprotein and contains many of the amino acid residues that have been experimentally characterized in terms of antibody binding and neutralization [3, 4]. However, sequence variation at an antibody-relevant residue does not by itself indicate altered neutralization, immune escape, vaccine failure, immune selection, or viral fitness. Therefore, interpreting sequence-level variation requires both careful documentation of the source of the data and clearly distinguishing between observed genomic variation and experimentally determined phenotypes. Sequence curation currently addresses several issues related to this issue. The Nextstrain platform offers open-source, reproducible phylogenetic surveillance workflows [5, 6], and the dengue workflow analyzed included an accession-level exclusion list that identified the focal records as duplicates of NC 001475 [7]. Additionally, a June 2026 bioRxiv preprint by de Moraes et al. (https://doi.org/10.64898/2026.06.12.731931) described methods for identifying matches between records across GenBank and GISAID and for removing source-specific sequence artifacts from viral phylogenomic datasets. While these examples highlight the importance of utilizing curated input data, they do not demonstrate that retaining a single representative of an identical-sequence group and removing the rest of that group results in the same site-level findings. Therefore, we compared these two analytical decisions using the same frozen dengue E-protein cohort. Our main analysis is a case study illustrating how cohort eligibility influences the amount of observed envelope-protein variation, not an estimate of the frequency of incorrect records in dengue databases. The separate global DENV3 screen identifies additional identity patterns for further review but does not establish that those records share the same mechanism or origin as the focal case. We used a frozen Bangladesh-associated dengue sequence cohort to assess whether representative-retaining deduplication, existing curated reference-aware exclusions, and provenance-informed cohort selection lead to the same E-protein variable-site classifications and candidate-prioritization decisions. We assessed exact sequence identity, complete-genome and archive context, an independently maintained Nextstrain exclusion snapshot, and two deduplication controls. The primary sequence-analysis endpoint was the set of E-protein positions classified as variable and potentially eligible for antibody-informed follow-up under each recordhandling rule, rather than the number of duplicates identified. The antibody-position enrichment analysis and historical sequence comparison were included as secondary, explicitly limited biological context.

## Methods

### Exact-identity provenance audit procedure

A separate computational process was developed to find exactly matching nucleotide sequences within public dengue records while keeping the identification of sequence-identity separate from the evaluation of the context of each record’s source.

#### Input records and normalization

The process accepts GenBank or FASTA inputs via Biopython. GenBank source features provide available record-level annotations; additionally, optional tab-separated accession metadata can be included for contextual review. Nucleotide sequences are converted to all-caps; spaces, tabs, carriage returns, and newlines are removed. The normalized sequence must be non-empty and consist only of A, C, G, T, or the supported IUPAC ambiguity symbols R, Y, S, W, K, M, B, D, H, V, and N. Sequences containing unsupported characters are rejected. For accession-level processing, terminal numeric accession version suffixes are removed when creating accession-base identifiers. Original record identifiers remain relevant to reporting and provenance interpretation.

#### Exact-identity detection and clustering

For each accepted nucleotide sequence, the process records its normalized sequence length and generates a SHA256 digest based upon its normalized nucleotide representation. Records are grouped together based on a joint key consisting of nucleotide sequence length and SHA256 digest. Only groups consisting of two or more records are reported as multi-record exact-identity clusters. Clusters are assigned sequential identifiers in the form EXACT 00001, EXACT 00002, etc., according to the deterministic grouping order of the implementation. Cluster summaries include membership counts and accession identifiers. Member-level outputs maintain the relationship between each input record and its assigned cluster. The SHA256 digest is utilized as a reproducible computational identity key. This process identifies matching normalized sequence representations; however, the resulting cluster does not independently establish biological sample identity or the reason for multiple archive representations.

#### Sequence-scope evidence classification

The process assigns a sequence-scope evidence label based upon nucleotide length and the record description. A record receives the tier1 explicit complete genome label if its sequence length is between 10,200 and 10,800 nucleotides, inclusive, and its description explicitly contains the phrase “complete genome”, regardless of letter case. A record in the same length range without that explicit description receives the genome length nonexplicit label. All other records receive the other sequence scope label. These labels describe available sequence-scope evidence. They are not classifications of provenance error, epidemiological independence, or record validity.

#### Metadata context and provenance interpretation

If available, the process records collection date information, geographic annotations, isolate/strain identifiers, host annotations, and accession-level metadata. A collection year is extracted if an eligible four-digit year is present in the appropriate metadata field. For each exact-identity cluster, the process provides a summary of the different collection years and geographic labels, earliest recorded collection year, corresponding accessions, and any RefSeq member accessions. These metadata summaries allow for contextual review of archive chronology and potential epidemiological/submission relationships. Metadata differences do not alter the underlying sequence-identity grouping. Therefore, complete-genome identity, record chronology, and submission context are evaluated independently prior to making a provenance-sensitive analytical decision. Exact E-region identity alone is not considered equivalent to complete-genome identity.

#### Validation and interpretation boundaries

The frozen standalone validation assessed the focal H001 complete-genome cluster and an independent plausible-linkage control. The H001 validation cluster consisted of 19 sequence-identical records: 18 Bangladesh accessions and the external comparison AY099336.1. Only the 18 Bangladesh records were excluded as per the prespecified H001 exclusion. The plausible-linkage control demonstrates that complete-sequence identity does not automatically warrant provenance-based exclusion. The process identifies sequence identity; it does not determine the mechanism responsible for producing identical records, establish whether independent/shared biological sampling occurred, or infer antibody phenotype, immune escape, adaptation, or viral fitness.

#### Reproducible implementation

The standalone audit implementation is provenance exact identity audit v1.py. Its record-level data, exact-cluster summaries, cluster-membership outputs, and audit summary provide a reproducible link between the analyzed input records and the reported sequence-identity relationships. The frozen validation results were verified prior to utilizing these relationships to interpret the focal H001 exclusion and subsequent analytical comparisons.

### Exact-identity workflow validation

The standalone exact-identity workflow was validated using a frozen 21-record validation fixture containing two previously identified sequence-identity clusters. Normalized nucleotide sequences were grouped by sequence length and SHA256 identity. Independent of downstream E-protein variation and antibody-enrichment results, recovery of the expected cluster memberships was evaluated. The H001 validation cluster consisted of 19 complete-genome records: 18 Bangladesh focal accessions and the older external accession AY099336.1. The 18 Bangladesh accessions, rather than all 19 cluster members, were the records subjected to the prespecified H001 exclusion. A separate plausible-linkage control contained two records, FJ850080.1 and FJ898447.1, with an identical 10,663-nt sequence. These records were evaluated separately from the 10,707nt H001 cluster. Exact identity was used for sequence grouping; contextual provenance classification remained a separate metadata-review step.

### Study design and frozen sequence cohort

This study was a retrospective computational analysis of publicly available dengue virus sequence records and associated archive metadata. No new biological samples were collected. The frozen high-quality E-protein cohort contained 426 accessions: 20 DENV1, 142 DENV2, 262 DENV3, and 2 DENV4. Residues were represented in serotype-specific E-protein coordinates and stored in a frozen long-format residue table containing accession, analysis serotype, collection year, E position, reference residue, observed amino-acid state, and callability. DENV2 and DENV3 were used for the primary comparative analyses because they had sufficient sequence representation. DENV1 was retained descriptively, and DENV4 was not analyzed further because only two sequences were available.

### DENV3 provenance and phylogenetic evaluation

The Bangladesh DENV3-2021 subset was examined at both the E-protein and complete-genome levels. Exact E-sequence haplotypes were identified, and representative sequences were placed in a global DENV3 E-protein phylogenetic context. The provenance-sensitive focal cluster was then evaluated using complete-record sequence length, exact nucleotide identity, archive dates, geography, isolate metadata where available, and complete-genome sequence hashes. The central comparison involved the Bangladesh focal records, the older external record AY099336.1, and the corresponding RefSeq representation NC 001475.2. Exact complete-genome identity was determined across the 10,707-nt records. The 18 Bangladesh focal records and AY099336.1 had zero nucleotide mismatches across the complete genome. The analysis establishes sequence identity and archive chronology only. It does not determine the mechanism that produced the identity.

### Prespecified H001 exclusion

A prespecified H001-exclusion scenario was evaluated for the 18 Bangladesh-attributed records whose 10,707-nt genomes matched the older external accession AY099336.1 exactly. This scenario tests how retaining versus excluding that sequence state changes the analytic cohort; genome identity and archive chronology do not establish that the reported biological specimens or collection metadata were erroneous. Ordinary Eprotein haplotype sharing, partial sequence identity, phylogenetic proximity, country, or collection year alone were not exclusion criteria. Application of this rule removed 18 accessions from the original 426-sequence cohort, producing an H001-excluded cohort of 408 sequences.

### Exact deduplication and H001-exclusion comparison

The initial comparison used four Bangladesh DENV3-2021 cohorts: the original 96 accessions, 47 accessions after whole-cohort exact-sequence deduplication, 79 after retaining one deterministic H001 representative (ON103302), and 78 after excluding the 18 focal Bangladesh records under the frozen provenance criterion. Two further comparators were evaluated using a saved external curation snapshot: NC 001475-specific accession exclusion (78 retained) and application of the entire static exclusion list (77 retained). These comparisons do not represent a rerun of the full Nextstrain phylogenetic workflow. For each cohort, E-protein positions were classified as variable when at least two callable amino-acid states were observed. The 13 positions that were variable in the original cohort but invariant after H001 exclusion were tracked across the deduplication controls.

Site-wise amino-acid frequency differences relative to the H001-excluded cohort were summarized using total variation distance (TVD). The comparisons distinguish changes in sequence multiplicity and amino-acid frequency distributions from changes in variable-site classification. For the external comparator, the accession exclusion set was constructed from the Nextstrain dengue phylogenetic/defaults/exclude.txt snapshot retrieved on October 21, 2026 [7]. Accession identifiers annotated ‘duplicate of NC 001475’ were selected without using frozen H001 labels. Their retained membership was then compared with the frozen H001-excluded cohort. A separate analysis applied every accession entry in the saved exclusion list to the same original 96-accession cohort. Variable-site classifications were recalculated from the frozen accession-level E-residue calls across 493 positions and checked against the frozen original and H001-excluded classifications. This analysis assesses static accession-list effects, not automated detection or provenance ground truth.

### E-protein variation analysis

For each analyzed serotype and time-defined subset, E-protein positions were classified as variable when at least two callable amino-acid states occurred among the included accessions. The primary provenance-impact comparison contrasted the original and H001-excluded Bangladesh DENV3-2021 subsets. Positions that disappeared after H001 exclusion were classified as H001-inclusion-dependent, whereas positions remaining variable after H001 exclusion were retained as variation in the H001-excluded cohort.

### Antibody-evidence atlas

Experimentally characterized antibody-relevant E-protein positions were assembled from the frozen literature evidence atlas [8–37]. To prevent temporal circularity, antibody positions used in the primary enrichment analysis were restricted to evidence available before the 2021 Bangladesh DENV3 sampling period. The final atlas contained 125 DENV2 E positions and 75 DENV3 E positions with pre-2021 literature support. Sequence variation at these positions was treated as antibody-relevant variation, not as evidence of phenotypic antibody escape.

### Positional enrichment analysis

Within each serotype, variable E positions in the analyzed cohort were compared with pre-2021 antibody-relevant positions. H001 exclusion affected DENV3 records only; the DENV2 cohort membership was unchanged. Observed overlap was compared with the corresponding E-protein positional universe. For DENV2 the universe contained 495 E positions, and for DENV3 the universe contained 493 positions. One-sided exact positional tests were used for the serotype-specific analyses. A serotype-stratified exact convolution was used for the combined DENV2/DENV3 result.

For each serotype *s*, the positional universe *N_s_* was the number of evaluated E-protein positions (495 for DENV2 and 493 for DENV3); *K_s_* was the number of positions in the pre-2021 antibody-evidence atlas (125 and 75); *n_s_* was the number of variable positions in the analysis cohorts (32 and 8); and *x_s_* was the observed overlap (10 and 2). Under the defined serotype-specific positional null, each subset of *n_s_* positions from *N_s_* was treated as equally likely. Accordingly, *X_s_ ∼* Hypergeometric(*N_s_, K_s_, n_s_*), the expected overlap was *n_s_K_s_/N_s_*, and the one-sided exact *p*-value was Pr(*X_s_ ≥ x_s_*). The combined analysis calculated Pr(*X*_DENV2_ + *X*_DENV3_ *≥* 12) from the exact convolution of the two serotype-specific hypergeometric distributions, assuming independent strata under this positional null; no pooled odds ratio was estimated.

Serotype-specific reported point odds ratios were sample cross-product odds ratios from 2 *×* 2 contingency tables of variable versus nonvariable positions and antibody-atlas membership. Their reported 95% confidence intervals were exact conditional intervals obtained by inversion of the noncentral hypergeometric distribution, as implemented using conditional estimation in `scipy.stats.contingency.odds ratio`. Thus, the point odds ratios and the confidence intervals use different estimators; the conditional odds-ratio estimates are not reported as the point estimates in the serotype-specific enrichment results. The unit of analysis is the E-protein position, not the accession or an independent infected individual. The positional null assumes exchangeability of E positions and does not model structural or functional dependence among positions, nor the nonrandom historical ascertainment of antibody-studied residues. These tests are descriptive evidence about positional overlap, not causal or phenotype-level tests. The enrichment analysis was prespecified as a positional analysis. Historical global variability was not used to modify or rescue statistical significance.

### Historical sequence-variation context

To determine whether H001-excluded Bangladesh variable positions or specific alternate amino-acid states had prior sequence precedent, public pre-2021 DENV2 and DENV3 E-protein backgrounds were examined. Exact-length E sequences were evaluated first. Coordinate compatibility with the frozen serotype reference sequences was verified before direct positional indexing. Non-exact-length records were then examined using mapped partial-E coverage for the small set of Bangladesh alternate states that were not found in the exact-length historical background.

Historical positional variability was calculated using distinct exact-length canonical E-protein amino-acid sequences as the primary unit. A position was classified as previously variable when at least two amino-acid states were observed, as recurrent when the non-major count was at least two, and as meeting the 1% threshold when the non-major frequency was at least 0.01. The frequency denominator was the number of distinct historical sequences evaluated at that position. Record-weighted frequencies were used only for sensitivity descriptions. These criteria do not establish independent evolutionary origins. Historical variation was used descriptively. It was not used as a covariate or post hoc adjustment to the antibody-position enrichment analysis.

### Global DENV3 E-identity screening and Tier-2 selection

The global DENV3 sequence-context assessment used the frozen set of 3,534 validated or rescued E-region records available to the study. Exact E-region sequence identity was used to group records, identifying 371 multi-accession E-identity clusters. This E-region screen was distinct from the Tier-1 strict complete-genome identity audit: identical E sequences alone were not regarded as evidence of complete-genome identity or of record error. The recorded Tier-2 screening script (`20a17b_prespecified_global_denv3_tier2_screen.py`) prioritized clusters using three binary indicators: records spanning at least two collection years, records representing at least two countries, and clusters containing at least five accessions. The raw priority score was the sum of these indicators (0–3). A single point was deducted, to a minimum of zero, if the recorded annotations identified any RefSeq member, patent context, explicit laboratory context, or passage context; multiple such annotations still produced only a one-point deduction. Clusters fully explained by a shared Tier-1 collision were ineligible. Among remaining eligible clusters, those with adjusted scores of at least two were selected for targeted manual review and available genome-length sequence comparison (25 clusters). This recorded prioritization rule identified review candidates, not proven provenance errors.

The subsequent isolate-label normalization repair corrected how nested isolatelabel pairs were represented after normalization; it was a reporting change rather than a change to the three scoring indicators or eligibility rules. The saved v1 and v2 selection tables retained the same 25 E-identity cluster hashes (25/25; none added or removed), with adjusted scores of two for 23 clusters and three for two clusters. Available genome-length records were subsequently compared within selected clusters to test whether E-region identity extended beyond E. Clusters lacking sufficient genome-length sequence evidence remained unresolved. Because selection was targeted rather than random and the underlying archive sampling was incomplete, these results cannot estimate provenance-error prevalence.

### Reproducibility

The frozen Bangladesh E-protein accession manifest includes 426 records; the specified H001 exclusion yields 408 retained accessions. Original and H001-excluded accession-level E-residue tables were checked for unchanged amino-acid calls in retained records. The temporally restricted antibody-position atlases contained 125 supported positions among 495 DENV2 positions and 75 among 493 DENV3 positions. The saved Nextstrain dengue exclusion file has SHA256 6a2e2a3645d3b53ca43b9fba864098e289bcfd57ff8afe1c732e15011b4123ff and matches the file at repository commit cca844c330aa9157bb3f34fc80cffab8935b7653 (committed December 18, 2024) [7]. A versioned staging package contains accession manifests, E-residue tables, position atlases, analysis scripts, supplementary materials, and check-sums. The checks establish agreement of frozen inputs and retained calls; they do not constitute an independent, clean-environment, end-to-end reproduction of every manuscript result.

### Use of Generative AI Assistance

ChatGPT (OpenAI) was utilized in this study for assistance debugging Python code, interpretation checks, and manuscript editing. This study utilized publicly available data and reproducible computational procedures. The authors are responsible for verifying the reported results, scientific interpretations, references, and final manuscript.

## Results

### Standalone exact-identity workflow recovers both validation cases

The standalone SHA256 audit recovered both expected clusters from the frozen 21-record validation fixture. The H001 cluster contained 19 records with identical 10,707-nt sequences, comprising the 18 Bangladesh focal accessions and AY099336.1. The independently characterized plausible-linkage control contained two records with identical 10,663-nt sequences. Both validation cases passed the prespecified exact-identity membership checks. The frozen contextual classification was provenance review required for H001 and consistent with plausible epidemiological linkage for the two-record control. These findings validate exact-identity recovery in the tested fixture, not the accuracy of automated provenance-error diagnosis. The workflow detects identical sequences; contextual interpretation requires independent metadata and archive-evidence review.

### A provenance-sensitive DENV3 cluster is complete-genome identical to an older external record

The original frozen cohort contained 426 high-quality E-protein sequences. Within the Bangladesh DENV3-2021 subset, 18 accessions formed a focal sequence cluster that required additional provenance evaluation. All 18 Bangladesh records were 10,707 nt long and were complete-genome identical to AY099336.1, with zero nucleotide mismatches across the entire genome. AY099336.1 is an older external record from Sri Lanka, whereas the Bangladesh records were labeled as collected in August 2021. The corresponding sequence is also represented by RefSeq NC 001475.2.

The shared complete-genome identity was reproduced during the final provenance audit. This establishes a provenance-sensitive identity cluster but does not establish why the records are identical. The 18 focal Bangladesh accessions were: ON103302, ON103390, ON111445, ON115021, ON115023, ON115025, ON115029, ON115182, ON115811, ON115814, ON115816, ON115818, ON116129, ON127418, ON127533, ON127537, ON127550, ON127552. These 18 records, together with the older external record AY099336.1, constitute the 19-member complete-genome exact-identity validation cluster. The 18 Bangladesh records were excluded under the prespecified H001 exclusion; AY099336.1 was an external comparison record and was not one of the 18 excluded Bangladesh accessions.

The complete-genome identity relationship is illustrated in Fig. 1.

**Fig. 1.**
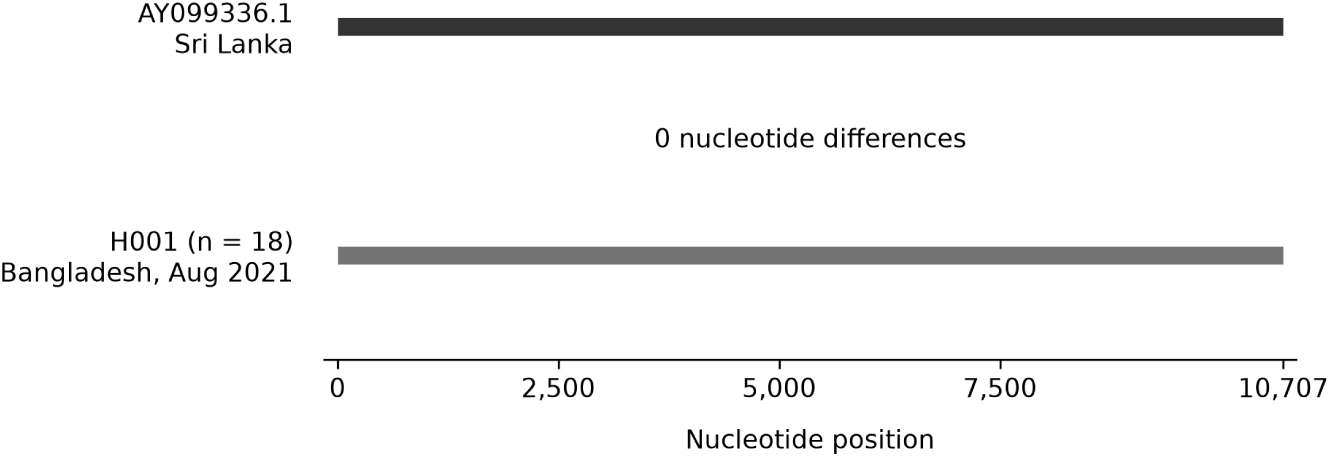
Complete-genome nucleotide identity between the Bangladesh H001 group and AY099336.1. Horizontal segments schematically represent the aligned 10,707-nt complete-genome sequence span of AY099336.1 and 18 Bangladesh-attributed DENV3 accessions (H001). The Bangladesh records have collection dates recorded as August 2021. No nucleotide differences were observed across the compared span. NC 001475.2 is the corresponding RefSeq representation and is not an additional member of the 19-record validation cluster. Exact identity does not establish biological sample independence or the mechanism of record identity. Accession-level details are provided in Online Resource 2

### H001 exclusion removes most of the apparent DENV3-2021 variable positions

Before H001 exclusion, the Bangladesh DENV3-2021 subset contained 96 accessions and 21 variable E-protein positions. After the 18 provenance-sensitive records were removed under the frozen criterion, 78 accessions remained and only 8 E positions remained variable. Thus, 13 of 21 originally variable positions (61.9%) disappeared after H001 exclusion. The positions lost after H001 exclusion were: E68, E81, E124, E132, E169, E219, E231, E270, E301, E377, E383, E404, E479. The positions remaining variable after H001 exclusion were: E221, E223, E224, E285, E303, E362, E375, E452. Notably, the previously observed variation at E270 and E383 disappeared completely after the provenance-sensitive records were removed. These results show that inclusion of the 18 records materially altered the apparent DENV3-2021 E-protein variation pattern. The primary cohort-level results are summarized in Table 1.

**Table 1.** Summary of the primary DENV3 provenance finding and analytical consequence.

| **Metric** | **Value** |
|---|---|
| Bangladesh DENV3-2021 accessions before H001 exclusion | 96 |
| Provenance-sensitive accessions excluded | 18 |
| Bangladesh DENV3-2021 accessions after H001 exclusion | 78 |
| Complete-genome length | 10707 |
| Mismatches between excluded Bangladesh records and AY099336.1 | 0 |
| Original variable DENV3-2021 E positions | 21 |
| Variable DENV3-2021 E positions after H001 exclusion | 8 |
| Variable E positions lost after H001 exclusion | 13 |
| Fraction of original variable sites removed | 61.9% |

The position-level amino-acid state counts for all 21 originally variable positions are given in Table 2.

**Table 2:** Position-level effect of H001 exclusion on Bangladesh DENV3-2021 E-protein variation.

| **E position** | **Reference** | **Original states** | **H001-excluded states** | **Excluded 18** | **After H001 exclusion** |
|---|---|---|---|---|---|
| 68 | I | V:78;I:18 | V:78 | I:18 | Lost |
| 81 | V | I:78;V:18 | I:78 | V:18 | Lost |
| 124 | P | L:78;P:18 | L:78 | P:18 | Lost |
| 132 | Y | H:78;Y:18 | H:78 | Y:18 | Lost |
| 169 | T | V:78;T:18 | V:78 | T:18 | Lost |
| 219 | A | T:78;A:18 | T:78 | A:18 | Lost |
| 221 | G | G:95;W:1 | G:77;W:1 | G:18 | Retained |
| 223 | T | I:70;T:26 | I:70;T:8 | T:18 | Retained |
| 224 | T | T:94;I:2 | T:76;I:2 | T:18 | Retained |
| 231 | R | K:78;R:18 | K:78 | R:18 | Lost |
| 270 | N | T:78;N:18 | T:78 | N:18 | Lost |
| 285 | L | L:95;P:1 | L:77;P:1 | L:18 | Retained |
| 301 | T | S:78;T:18 | S:78 | T:18 | Lost |
| 303 | T | A:77;T:18;V:1 | A:77;V:1 | T:18 | Retained |
| 362 | P | L:77;P:18;V:1 | L:77;V:1 | P:18 | Retained |
| 375 | N | N:95;T:1 | N:77;T:1 | N:18 | Retained |
| 377 | V | I:78;V:18 | I:78 | V:18 | Lost |
| 383 | N | K:78;N:18 | K:78 | N:18 | Lost |
| 404 | E | A:78;E:18 | A:78 | E:18 | Lost |
| 452 | V | I:77;V:18;T:1 | I:77;T:1 | V:18 | Retained |
| 479 | A | V:78;A:18 | V:78 | A:18 | Lost |

Inspection of the frozen accession-level state counts showed complete segregation at all 13 E-protein positions that became invariant following exclusion of the 18 Bangladesh-attributed H001 records. At each position, all 18 H001 records carried one amino-acid state, whereas all 78 retained records carried a different state (13/13 positions; Table 2). Retaining even one H001 representative therefore preserved two observed states at these positions, explaining why representative-retaining deduplication preserved all 21 variable-site classifications. Of the eight positions remaining variable after H001 exclusion, six had a minor amino-acid state observed in only one accession. The other two were E223 (minor-state count, eight) and E224 (minor-state count, two). The four-cohort variable-site classification matrix is provided as Online Resource 4.

### Representative-retaining deduplication and curated cluster exclusion produce different variable-site calls

In the original Bangladesh DENV3-2021 cohort, 96 accessions contained 21 variable E-protein positions. Ordinary whole-cohort exact full-sequence deduplication reduced the cohort to 47 accessions but retained all 21 variable positions. Collapsing only the 18 Bangladesh H001 records to one deterministic representative yielded 79 accessions and likewise retained all 21 variable positions. In contrast, the prespecified H001 exclusion removed the 18 focal records, leaving 78 accessions and eight variable E positions. All 13 positions lost after H001 exclusion remained variable in both representative-retaining deduplication controls. Thus, reducing the number of identical sequence records while retaining one H001 representative did not reproduce the variable-site classification obtained when the complete focal group was excluded. The independently curated Nextstrain exclusion entries annotated as duplicates of NC 001475 identified exactly the same 18 Bangladesh accessions as the prespecified H001 exclusion. All 96 original accessions had identical retained/excluded membership under these two decisions. Both produced 78 retained accessions and eight variable E positions. The entire saved Nextstrain exclusion list additionally excluded OP741243, annotated as a duplicate of OK079104, leaving 77 accessions. Independent recalculation across 493 E positions identified eight variable sites and no changed variable-site classifications relative to the 78-accession cohort.

At the 13 provenance-dependent positions, the H001 sequence contributes an amino-acid state absent from the other 78 records. Keeping one representative therefore preserves the classification of these positions as variable, whereas excluding the complete group removes that additional state. This is a cohort-eligibility effect rather than an effect attributable to duplicate multiplicity alone. The effects of the six record-handling strategies on accession retention and variable-site classification are summarized in Fig. 2. Frequency distortion was reduced by deduplication. Mean site-wise TVD relative to the H001-excluded cohort was 0.00645 for the original cohort, 0.000853 for the whole-cohort deduplicated cohort, and 0.000436 for the H001 single-representative sensitivity cohort. A position-resolved heatmap shows these differences at the 21 positions variable in the original cohort (Fig. 3). The maximum sitewise TVD decreased from 0.1875 for the original cohort to 0.0297 after whole-cohort deduplication. Exact deduplication substantially reduced differences in state frequencies relative to the H001-excluded cohort but retained all 13 focal variable-site calls. The independently documented reference-specific exclusions reproduced the H001-excluded cohort membership, and the additional static-list exclusion did not change site-level variability. The comparison thus separates effects of sequence multiplicity from decisions about which sequence states belong in the analytical cohort.

**Fig. 2.**
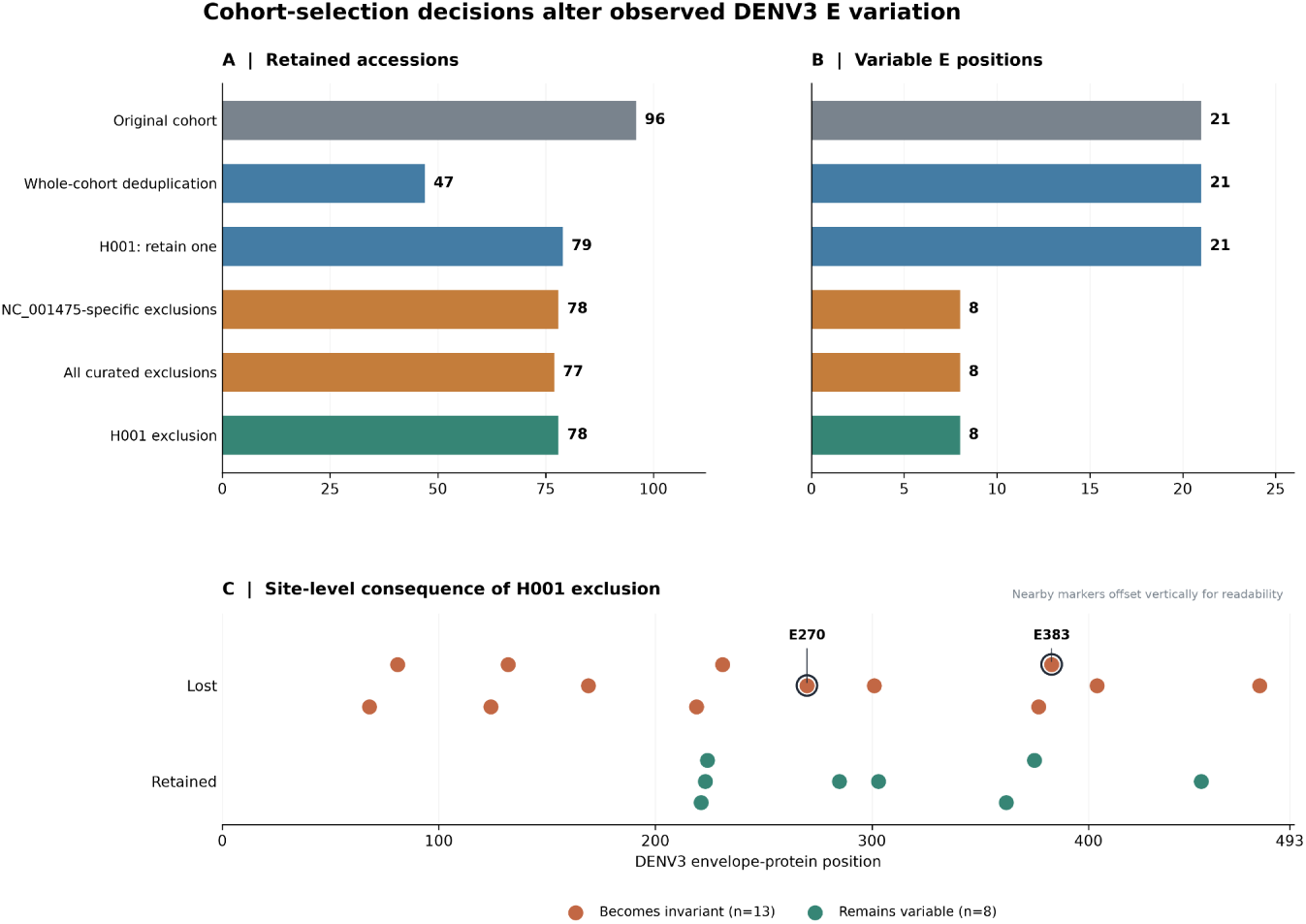
Effects of sequence-record handling on DENV3 E-protein variation. Panels A and B compare accession retention and numbers of variable E-protein positions under six analytical strategies applied to the same frozen Bangladesh DENV3-2021 cohort. The two representative-retaining deduplication controls preserve one member of the H001 group, whereas the focal H001-exclusion and Nextstrainlist scenarios remove the relevant accession records entirely. Panel C shows the 13 positions becoming invariant and the eight positions remaining variable after exclusion of the 18 Bangladesh-attributed H001 records. Horizontal locations represent the actual E-protein coordinates; nearby markers are displaced vertically for readability. E270 and E383 are highlighted because of previously published antibody-related evidence. The Nextstrain comparisons apply a saved accession-exclusion list rather than rerunning the entire phylogenetic workflow. The analytical strategies are not independent biological replicates, and no antigenic phenotype is inferred

**Fig. 3.**
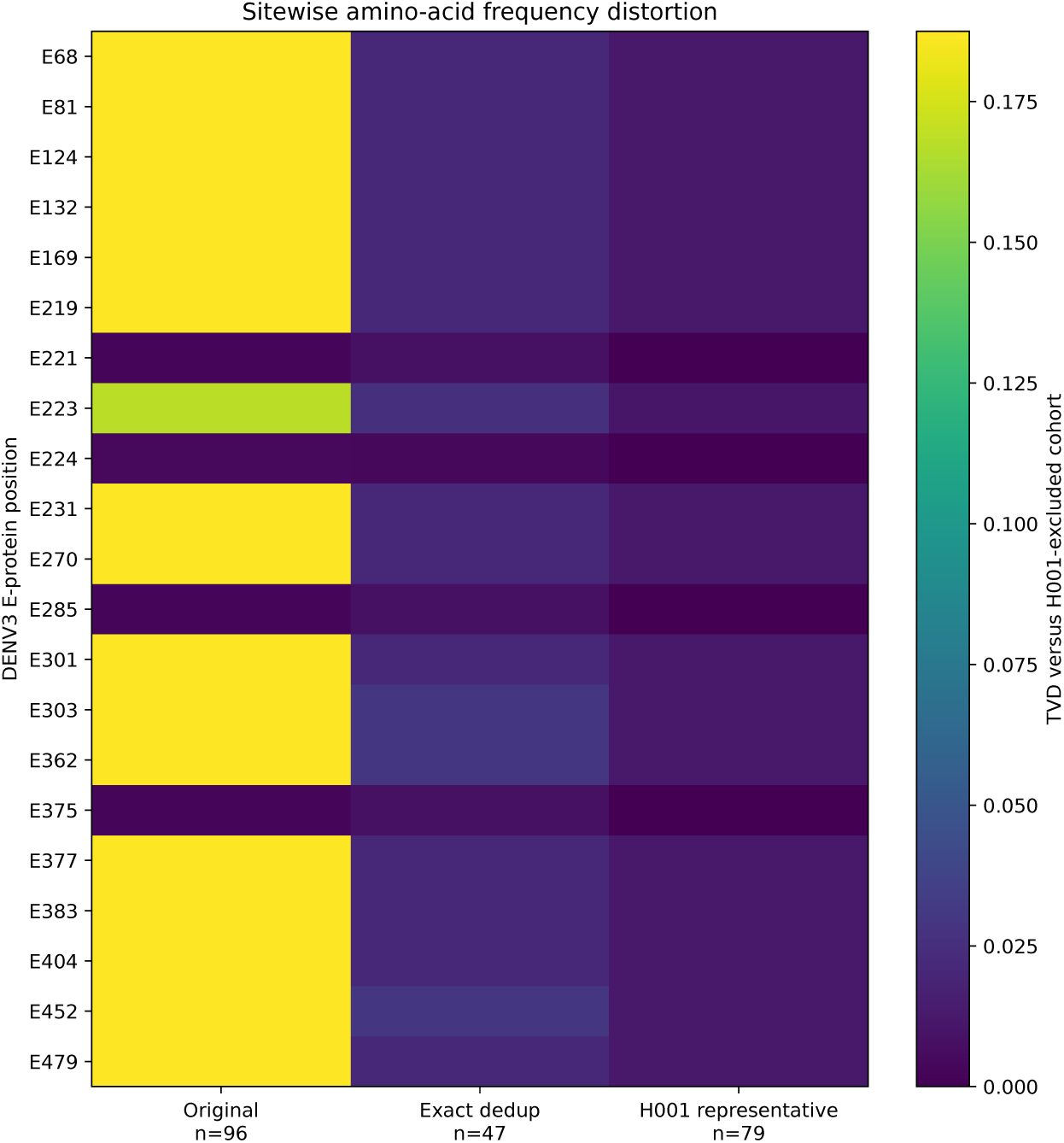
Sitewise amino-acid frequency distortion under three DENV3 sequence-record handling strategies. Total variation distance (TVD) at each E-protein position was measured relative to the H001-excluded cohort (78 accessions). The heatmap displays only the 21 E-protein positions variable in the original 96-accession cohort. Columns represent the original cohort (96 accessions), whole-cohort exact-sequence deduplication (47 accessions), and retention of one H001 representative (79 accessions). The cohort-wide mean TVDs reported in the Results were calculated across all 493 evaluated E-protein positions, not only the 21 displayed here. Higher TVD indicates a larger difference in the observed amino-acid state frequencies, not independent mutation events or experimentally measured antigenic effects

### H001 exclusion changes antibody-informed variant prioritization

The provenance-dependent E-protein positions were examined for their consequences for research prioritization using the frozen pre-2021 antibody-evidence atlas. DENV3 E270 and E383 illustrate this consequence. Both positions were variable in the original Bangladesh DENV3-2021 dataset and remained variable after ordinary whole-cohort exact-sequence deduplication. Both positions also remained variable when the 18 focal H001 accessions were collapsed to one representative. Historical antibody evidence provides context for these positions, but the evidence differs in kind. For E270, the 5J7 escape-selection study reported a lysine insertion between Q269 and N270 [26]; it did not establish a neutralization effect for the N270T substitution observed in the original Bangladesh cohort. For E383, the frozen atlas includes structural and functional evidence involving antibodies 4E11 and 8A1 [24, 29]. Thus, E270 and E383 fulfilled the prespecified combination of observed sequence variation and historical antibody-related evidence, without implying that their observed substitutions had experimentally demonstrated effects. After exclusion of the 18 provenance-sensitive records, both positions became invariant in the H001-excluded Bangladesh DENV3-2021 subset. Consequently, E270 and E383 no longer qualified for prioritization based on variation in the H001-excluded cohort. In contrast, E223 and E224 remained variable after H001 exclusion (Table 2; Online Resource 1). Both positions were reported among the modeled 5J7 antibody-interface residues in the pre-2021 cryo-EM study [33]. These structural observations do not establish altered 5J7 binding or neutralization caused by the amino-acid states observed in the Bangladesh sequences.

This demonstrates that provenance evaluation can alter which E-protein positions are selected for downstream hypothesis generation, even when routine exact-sequence deduplication leaves their variable-site status unchanged. The analysis concerns computational prioritization criteria only. It does not demonstrate altered antibody binding, neutralization, immune escape, vaccine performance, or viral fitness at either position. The accession-level support for the eight retained variable positions is summarized in Fig. 4.

**Fig. 4.**
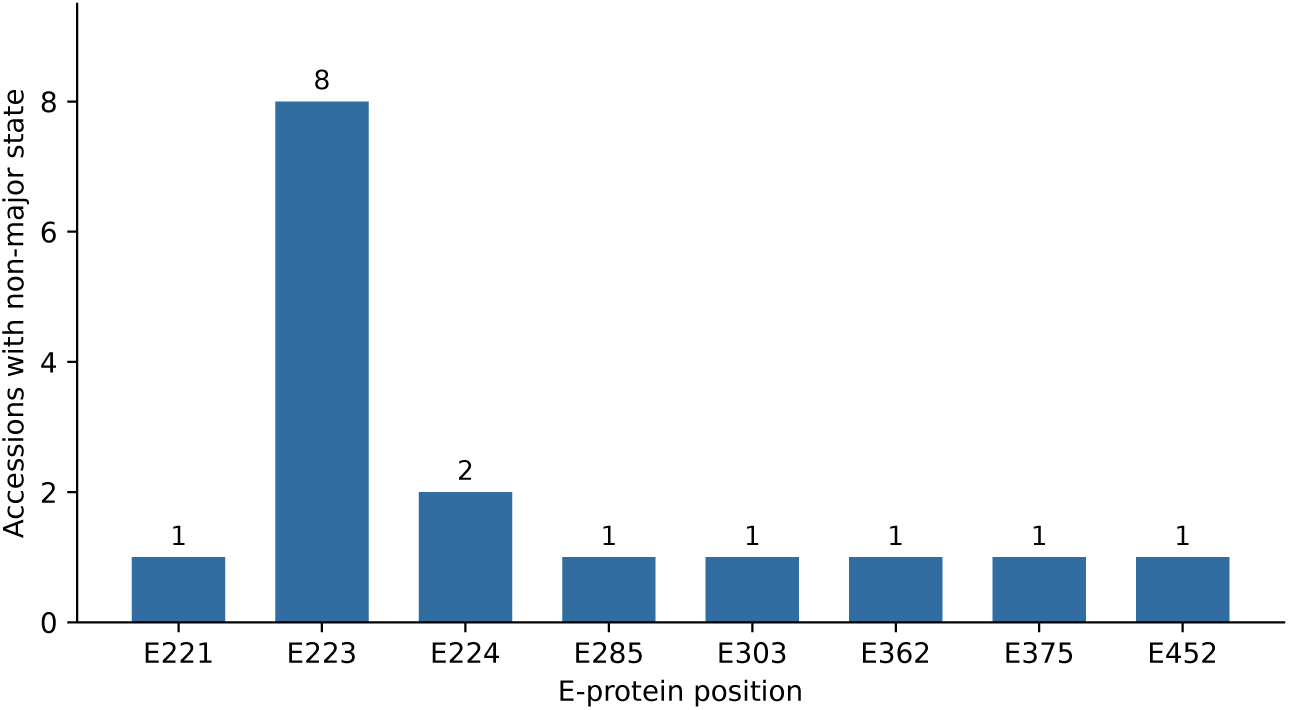
Minor amino-acid-state accession counts among the eight variable DENV3 E-protein positions remaining after H001 exclusion. Counts reflect the number of records carrying a non-major amino-acid state among the 78 retained Bangladesh DENV3-2021 accessions. Six positions are supported by a single non-major-state accession, E224 by two, and E223 by eight. Accession counts do not estimate independent mutational origins or demonstrate experimentally measured antigenic effects

### Exact complete-genome provenance-review patterns recur in the examined global DENV3 archive

To assess whether the focal H001 pattern was the only provenance-review-worthy exact-identity event in the examined data, we reviewed strict complete-genome collisions in the available global DENV3 corpus. The Tier-1 audit identified 20 exact complete-genome collision clusters. Contextual review classified 16 as consistent with plausible epidemiological linkage, three as requiring provenance review, and one as clearly laboratory-derived. The three provenance-review-required clusters comprised H001 and two distinct Puerto Rico multi-year collision clusters. Thus, complete-genome identity patterns requiring provenance review were not confined to the focal Bangladesh event. A separate Tier-2 E-region identity screen identified 371 multi-accession E-sequence collision clusters among 3,534 validated or rescued DENV3 E records. E-region identity was treated as screening evidence rather than proof of complete-genome identity. Of 25 clusters selected under the recorded Tier-2 screening rule, 13 contained available genome-length sequences that differed outside E, three contained exact-genome subclusters already represented in Tier 1, and eight remained unresolved because fewer than two genome-length records were available. The remaining selected cluster yielded an additional exact 10,640-nt sequence pair, ON890819 and ON907582. Both records shared a collection date of 8 July 2019, country China, normalized isolate identifier, and study/submission context. This pair was therefore considered consistent with non-independent archive representation of the same isolate for event counting. The process by which the two accessions entered the archive was not established. These findings demonstrate recurrence of strict complete-genome identity patterns warranting provenance review within the examined global DENV3 corpus. They do not establish that all collisions represent archive errors or share a common mechanism. The analysis was restricted to the available global DENV3 corpus and was not designed to estimate the prevalence of provenance errors. Tier-2 E-region identity alone was not used to expand the set of complete-genome provenance-review events. The global DENV3 identity-cluster review did not result in any additional exclusions from the frozen Bangladesh analytical cohort.

### DENV2 and H001-excluded DENV3 variation is not significantly enriched at antibody-relevant positions

The unchanged DENV2 cohort contained 32 variable E positions, of which 10 overlapped the 125 pre-2021 antibody-relevant positions. The expected overlap under the positional null was 8.08, corresponding to a fold enrichment of 1.24 and a one-sided exact p-value of 0.26896. The H001-excluded DENV3 cohort contained eight variable E positions, of which 2 overlapped the 75 pre-2021 antibody-relevant positions. The expected overlap was 1.22, corresponding to a fold enrichment of 1.64 and a one-sided exact p-value of 0.349767. In the serotype-stratified combined analysis, 12 overlaps were observed versus 9.30 expected (one-sided exact convolution p=0.19446794). The exact conditional 95% odds-ratio confidence intervals were 0.56 to 3.13 for DENV2 and 0.18 to 10.77 for DENV3. Both intervals include the null odds ratio of 1. In particular, the wide DENV3 interval reflects limited precision from the small number of variable positions in the analysis cohorts. These results do not establish the absence of antibody-position enrichment; rather, statistically significant enrichment was not detected in the examined data. None of these analyses provided statistically significant evidence of positional enrichment. Observed and expected positional overlaps, together with their statistical uncertainty, are illustrated in Fig. 5.

**Fig. 5.**
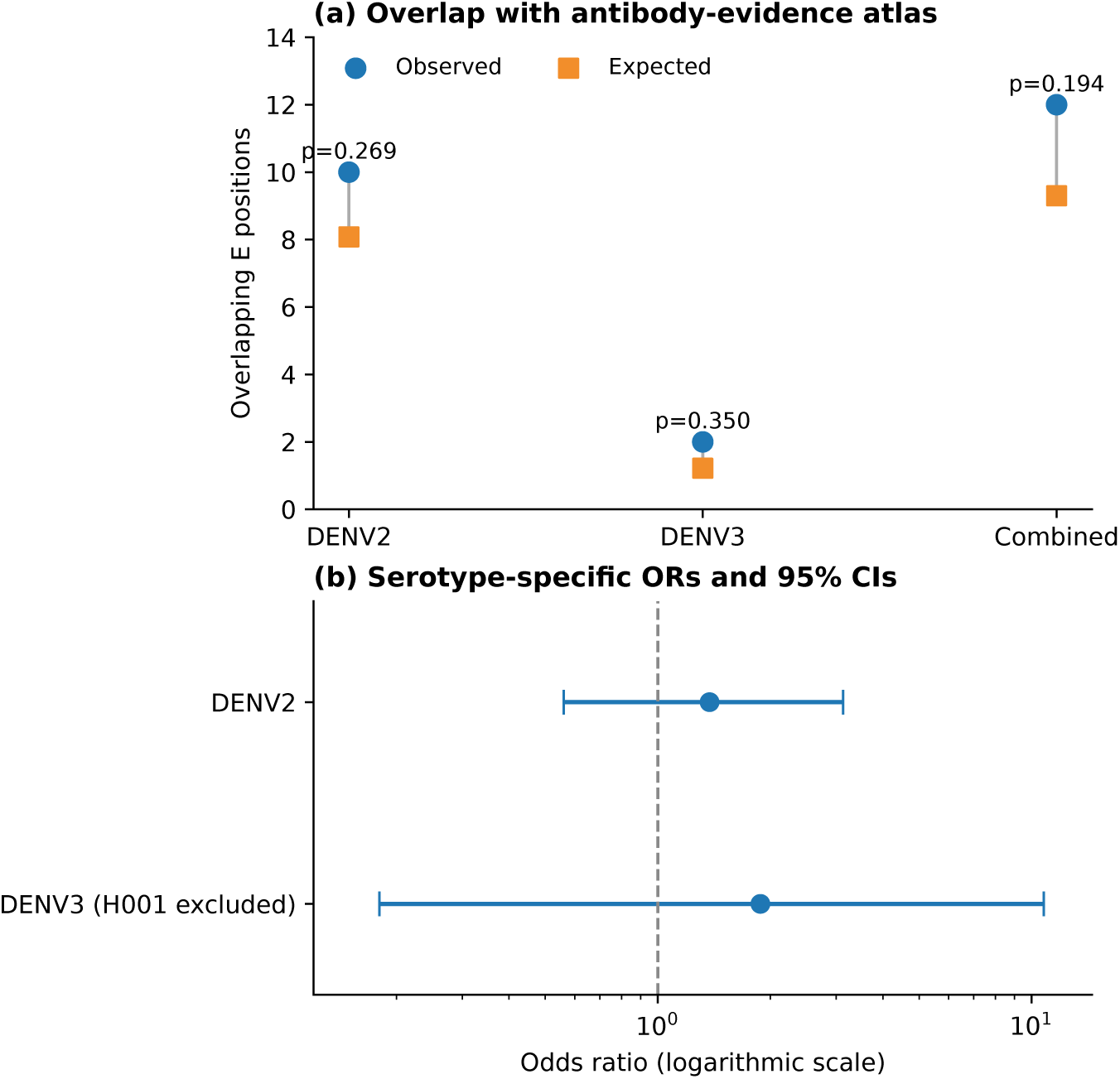
Antibody-relevant positional overlap and uncertainty in the analyzed DENV2 and H001-excluded DENV3 cohorts. (a) Observed overlaps with the frozen pre-2021 antibody-position atlas are shown alongside the expectations under the specified serotype-specific positional null; the combined comparison uses a serotype-stratified exact convolution. (b) Serotype-specific sample cross-product odds-ratio point estimates and exact conditional 95% confidence intervals are shown on a logarithmic scale, with the null odds ratio of one marked by a dashed line. Neither serotype-specific test nor the stratified combined analysis detected statistically significant positional enrichment. No combined odds ratio was estimated. Positions, not accessions or independent evolutionary events, are the units of the positional tests

The serotype-specific and combined positional tests are summarized in Table 3.

**Table 3.**
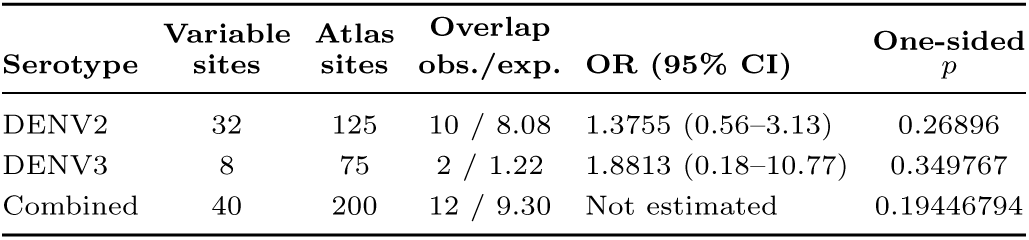
Antibody-relevant positional overlap in the DENV2 and H001-excluded DENV3 cohorts. Overlap expectations follow the defined positional null; odds ratios (ORs) are sample cross-product estimates with exact conditional 95% confidence intervals (CIs). The combined row gives the serotype-stratified exact-convolution *p*-value; a pooled OR was not estimated.

### Most variable positions in the analysis cohorts had historical sequence precedent

All 32 DENV2 variable positions had shown amino-acid variation in the examined public pre-2021 background. Of the 8 H001-excluded DENV3 variable positions, 6 had prior variation in the historical background. At the specific amino-acid-state level, six Bangladesh non-major states were not observed in either the exact-length or mapped partial pre-2021 sequence background: E235D, E307T, E354I, E221W, E285P, E362V. Five of these six observations were singletons. DENV2 E354I occurred in seven Bangladesh accessions representing six distinct E nucleotide sequences, but all seven accessions belonged to one submission/provenance context. Accordingly, the historical-background analysis provides descriptive context for the H001-excluded Bangladesh variation but does not establish independent emergence, adaptive evolution, altered antibody phenotype, or increased viral fitness. Historical position and state summaries are reported in Table 4.

**Table 4.**
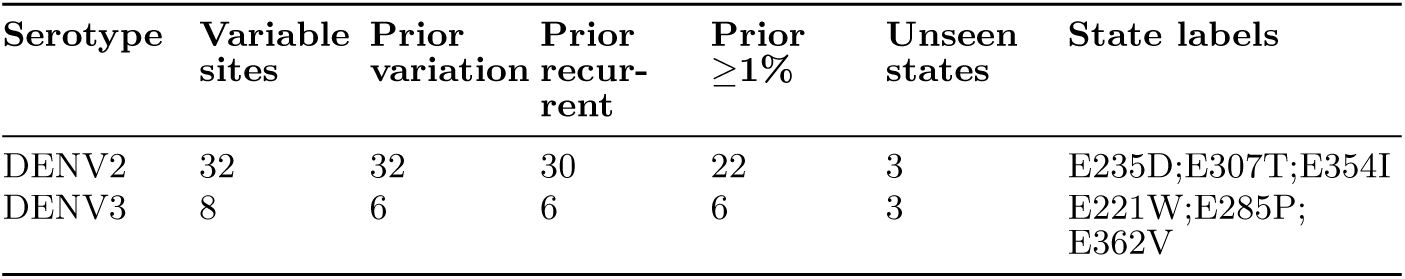
Historical pre-2021 sequence context for H001-excluded Bangladesh E-protein variation.

Online Resource 3 (ESM 3.csv) provides the six Bangladesh non-major amino-acid states not observed in the examined pre-2021 backgrounds, with supporting accession-level context. The underlying records and interpretations are unchanged.

### Published accession-label appearance and limits of sequence-level reuse evidence

A source-bounded review examined whether the 18 focal Bangladesh DENV3 accession identifiers had documented downstream visibility. Sources included accession-linked PubMed and PMC searches, substantive GenBank references, non-self NCBI public-data records, DataCite and Zenodo metadata, and relevant published phylogenetic figures. The database-based screening found no verified exact accession mentions within the successfully examined sources. However, 18 search-source failures limited the coverage of that screening. Its negative result therefore cannot be interpreted as evidence of no downstream use. Inspection of Figure 6 in Hasan and Islam (2024) [38] identified all 18 focal H001 accession identifiers in a published phylogenetic figure labeled as a DENV1 E-gene analysis. The corresponding GenBank accession records examined here are annotated as DENV3. This observation establishes accession-label appearance in a published figure, but does not independently establish inclusion of the corresponding nucleotide sequences in the original phylogenetic alignment or tree reconstruction. The underlying analysis inputs were not verified in this review. We therefore distinguish documented published accession-label appearance from verified sequence-level analytical reuse. The figure-label discrepancy does not establish a specific analytical error, a shared specimen origin, or the mechanism responsible for the H001 complete-genome identity.

### All focal records remained retrievable in the dated archive snapshot

A dated NCBI archive-status assessment was performed for the 18 focal Bangladesh DENV3 accessions. At the October 20, 2026 snapshot (23:39:31 UTC), all 18 records were retrievable. All 18 retrieved sequences matched the frozen 10,707-nt H001 sequence. No retrieved sequence differed from the frozen H001 sequence, and none of the 18 accessions was unresolved. No suppression, withdrawal, replacement, or correction status terms were identified in the examined records. No retrieval errors were reported. This snapshot establishes the observable archive state at the specified retrieval time. It does not demonstrate whether the records were reviewed by archive curators, whether the submitted metadata were independently validated, or why the records remained available. No change to the prespecified H001 exclusion or the scientific classification of the focal records resulted from the archive-status assessment.

## Discussion

The primary finding of this study is related to analytical cohort eligibility, rather than a demonstrated dengue phenotype through experimentation. There were 18 Bangladesh-associated DENV3 records that were completely identical to an older external sequence at the nucleotide level across all 10,707 nucleotides. When the H001 sequence state was included, there were 21 variable E positions in the original 96-accession cohort. However, when all 18 focal records were excluded, there were only eight variable positions among the 78 accessions. Therefore, 13 of the 21 originally variable positions (61.9%) were dependent upon the inclusion of that sequence group.

While reducing the multiplicity of that sequence group without removing every representative did not eliminate those site-level differences, all 13 H001-exclusion-dependent positions demonstrated complete state separation between the 18 focal and 78 retained records. Thus, retaining a single H001 representative allowed for the preservation of the second observed state and the classification of each of these positions as variable; only excluding all 18 focal records removed that state from the analytical cohort. This explains why exact-sequence deduplication alone did not produce the same result as H001 exclusion. Nonetheless, six of the eight positions remaining variable after exclusion had a minor state supported by only one accession; therefore, without read-level verification, sequencing, assembly, or alignment artifacts remain plausible explanations for individual singleton calls. The observed identity pattern does not provide evidence of invalid biological specimens or an antibody-related phenotype. This study illustrates why archive provenance should be evaluated prior to interpreting unusual sequence patterns as biological evolution.

Similar studies have shown that the consequences of database quality decisions vary depending on the downstream endpoint. Chen et al. studied how different types of duplicates affect summary measurements [2]; the 2026 bioRxiv preprint by de Moraes et al. described cross-repository reconciliation and laboratory-associated sequence artifacts using chikungunya data; and Hunt et al. quantified the effects of systematic SARS-CoV-2 sequence reconstruction errors on phylogenetic interpretation [39]. Our measured endpoint is narrower: which DENV3 E-protein positions are called variable and consequently considered for antibody-informed computational prioritization. We did not estimate changes in evolutionary rates, phylogenetic introductions, or experimental antigenic phenotypes. Original-study provenance helps interpret the retained DENV3 variation. Rahim et al. reported E223 variation and T224I changes in clinical DENV3 E-region sequences from Dhaka [40]. Across the full deposited range OQ826841–OQ827019 (179 accession records from multiple years and serotypes), 40 accessions (OQ826941–OQ826980) occur in our original 96-record DENV3-2021 cohort. All 40 are retained after exclusion of H001 (40/78; 51.3%). None belongs to the 18 Bangladesh-attributed H001 accessions, which have ON-prefix identifiers. Therefore, the Rahim study documents the originating clinical sequencing study and previously reported E223/E224 changes; these overlapping accessions do not constitute independent external validation. Agreement with the published study also does not independently establish the raw-read quality of individual amino-acid calls.

A separate published Dhaka DENV3 study by Jahan et al. reported E-protein substitutions G221W and L285P, along with T223I, among 17 representative E-gene sequences from 2021 [41]. These published residue-level observations provide contextual precedent for E221 and E285, two of the six singleton-minor-state positions in our H001-excluded cohort. We did not verify that the corresponding Jahan accessions were disjoint from our frozen cohort, nor did we compare the original read-level evidence. Therefore, the study cannot be treated as independently validated support for the particular accession-level calls in this analysis. Existing Nextstrain dengue curation had already identified the focal accessions for exclusion [7]; therefore, we do not claim first discovery or first exclusion. The specific contribution of this worked comparison is to quantify why an identical-sequence group and its cohort membership cannot be treated as interchangeable with simple sequence redundancy: representative retention preserves all 13 H001-specific states, whereas complete exclusion removes them from the 2021 Bangladesh analytic cohort. The resulting variable-site and antibody-informed prioritization differences are reproducible conditional on these cohort decisions. Exact identity and chronology do not establish why the records are identical or prove that their collection metadata are erroneous.

The antibody-position overlap analysis also changes the emphasis of the study. Although DENV2 and DENV3 variable positions showed modest numerical excesses at literature-defined antibody-relevant positions, neither serotype-specific test nor the combined stratified analysis was statistically significant. Therefore, the analyzed positional data do not support a claim that Bangladesh E-protein variation is preferentially concentrated at experimentally characterized antibody-relevant positions. The historical sequence analysis provides a further constraint on interpretation. Most variable positions in the analysis cohorts had already varied in public pre-2021 sequence backgrounds. Six particular Bangladesh alternate amino-acid states were not observed in the historical backgrounds examined, but five were singletons and the only recurrent example, DENV2 E354I, was confined to one provenance context. These observations are suitable for descriptive reporting but are insufficient to establish independent emergence, positive selection, antibody escape, or altered viral fitness.

These findings have implications for interpreting public dengue genomic surveillance data. Whether an E-protein state appears in deposited records is distinct from whether that state remains represented among records eligible for a specified surveillance cohort. Here, the distinction changed variable-site classification at 13 E positions and the eligibility of E270 and E383 for variation-based antibody-informed prioritization. Therefore, reporting an exact-deduplication rule alone is insufficient when a provenance-sensitive sequence group carries a state absent from the retained cohort. Explicit accession-level eligibility criteria and side-by-side site classifications make the resulting interpretations more transparent and reproducible.

This is a recommendation for computational interpretation, not evidence of altered viral phenotype, epidemic transmission, or incorrect specimen metadata. The examined Nextstrain dengue workflow provides important prior curation evidence [7]. Its saved exclusion file explicitly lists the 18 focal Bangladesh accessions and AY099336 as duplicates of NC 001475. Applying those reference-specific annotations to our frozen 96-accession cohort reproduced membership of the 78-accession H001-excluded cohort exactly. Applying the full static exclusion list also removed OP741243, without changing the eight remaining variable E positions. These results do not imply that Nextstrain performed independent provenance adjudication or that its complete phylogenetic workflow was reproduced here. They demonstrate agreement with existing curation while clarifying which duplicate-handling semantics change this cohort’s inferred diversity. This study has several limitations. First, it is based on public archive records and therefore inherits limitations of submitted metadata and sampling coverage. Second, the H001 exclusion is a provenance-informed sensitivity scenario, not independent proof of archive error or invalid specimens; other forms of sequence and sampling non-independence may remain. Third, the antibody analysis is positional and literature-based; positions with extensive experimental study may differ systematically from less studied positions. Fourth, sequence variation alone cannot establish antibody neutralization phenotype, vaccine performance, disease severity, ADE, immune selection, or experimental viral fitness. Finally, absence of a specific amino-acid state from the examined historical background should be interpreted as absence from that dataset, not proof that the state had never occurred globally. Sequence-record counts must also be distinguished from independent evolutionary events. Forty of the 78 retained DENV3-2021 accessions originate from one clinical sequencing study, and related sequences may share amino-acid states through common ancestry. Accordingly, the eight records carrying the minor E223 threonine state and the two carrying the minor E224 isoleucine state must not be interpreted as eight and two independent mutational origins. We did not estimate the number of independent mutation events, and the singleton-supported state calls have not been independently verified using raw-read evidence in this accession-based analysis.

## Conclusion

This computational study shows that representative-retaining exact-sequence deduplication and complete exclusion of an identical-sequence group can produce different variable-site classifications. Within the Bangladesh DENV3-2021 cohort, both representative-retaining controls preserved 21 variable E positions, whereas exclusion of all 18 focal H001 accessions yielded eight. Existing Nextstrain reference-specific curation selected the same 18 accessions, and its full saved exclusion list removed one additional record without changing the eight-position result. This distinction also affected downstream research prioritization. DENV3 E270 and E383, which had independent historical antibody-related evidence, remained variable after both deduplication controls but became invariant in the H001-excluded cohort. Thus, the difference between deduplication and provenance validation extended beyond record counts to the selection of candidate positions for biological follow-up. The standalone exact-identity workflow provides a reproducible computational means of detecting sequence-identity clusters, while the plausible-linkage validation control demonstrates why identity alone cannot determine provenance. Additional complete-genome identity patterns requiring contextual review were observed in the examined global DENV3 corpus, without supporting an archive-wide prevalence estimate. Together, these findings support evaluating exact-sequence identity alongside archive metadata and making cohort-eligibility choices explicit before interpreting dengue E-protein variation or nominating positions for biological investigation. Unlike representative-retaining deduplication, complete exclusion can change which amino-acid states remain in an analysis cohort. The conclusions concern reproducibility of sequence-based genomic surveillance and research prioritization; they do not establish a provenance mechanism or experimentally measured antigenic or fitness effects.

## Supporting information

Supplementary Figure S1 - DENV3 E-protein amino-acid composition

Supplementary Table S1 - Dengue accession provenance records

Supplementary Table S2 - Historically unseen amino-acid states

Supplementary Figure S2 - DENV3 variable-site classification matrix

## Data and code availability

The dengue nucleotide sequences analyzed in this study were obtained from public GenBank records. The accession manifests and supplementary tables document cohort membership and derived E-protein states. A versioned research package contains the frozen sequence inputs, accession manifests, antibody-position atlases, analysis scripts, selected outputs, figures, and SHA256 checksums. Two core analytical workflows were validated using reproduce core.py; the package does not independently reproduce every manuscript figure or table. The Nextstrain exclusion list is cited by its pinned Git revision and SHA256 checksum [7] and is not redistributed. Original analysis software is licensed under MIT; third-party GenBank and Nextstrain materials retain their own applicable rights. The research package is not yet in a public archive; materials can be requested from the corresponding author for editorial and peer-review assessment. No new biological specimens or viral sequences were generated.

## Statements and Declarations

## Ethics approval and consent to participate

Not applicable. The study analyzed publicly available viral nucleotide sequences and archival metadata; no participants were recruited and no new biological samples were collected.

## Supplementary information

**Online Resource 1 (ESM 1.pdf).** Amino-acid state composition at four DENV3 envelope-protein positions before and after H001 exclusion.

Panels show E223, E224, E270, and E383 in the original 96-accession cohort and the 78-accession H001-excluded cohort. E223 and E224 remain variable after exclusion, whereas E270 and E383 become invariant. Counts represent sequence records, not independent evolutionary events or experimentally measured antibody effects.

**Online Resource 2 (ESM 2.csv).** Archive metadata and complete-genome provenance evidence for the 18 Bangladesh records, AY099336.1, and NC 001475.2.

**Online Resource 3 (ESM 3.csv).** Bangladesh amino-acid states not observed in the examined pre-2021 exact-length or mapped partial E-protein backgrounds.

**Online Resource 4 (ESM 4.pdf).** Variable-site classification matrix for the 21 originally variable Bangladesh DENV3-2021 E-protein positions across the original, exact-deduplicated, single-H001-representative, and H001-excluded cohorts. V denotes a variable position, and a dash denotes an invariant position. The matrix does not represent independent evolutionary events.

