## Supplementary Figure S1 - DENV3 E-protein amino-acid composition for "Provenance-aware handling of identical dengue genome records alters inferred envelope-protein variation in Bangladesh DENV-3"

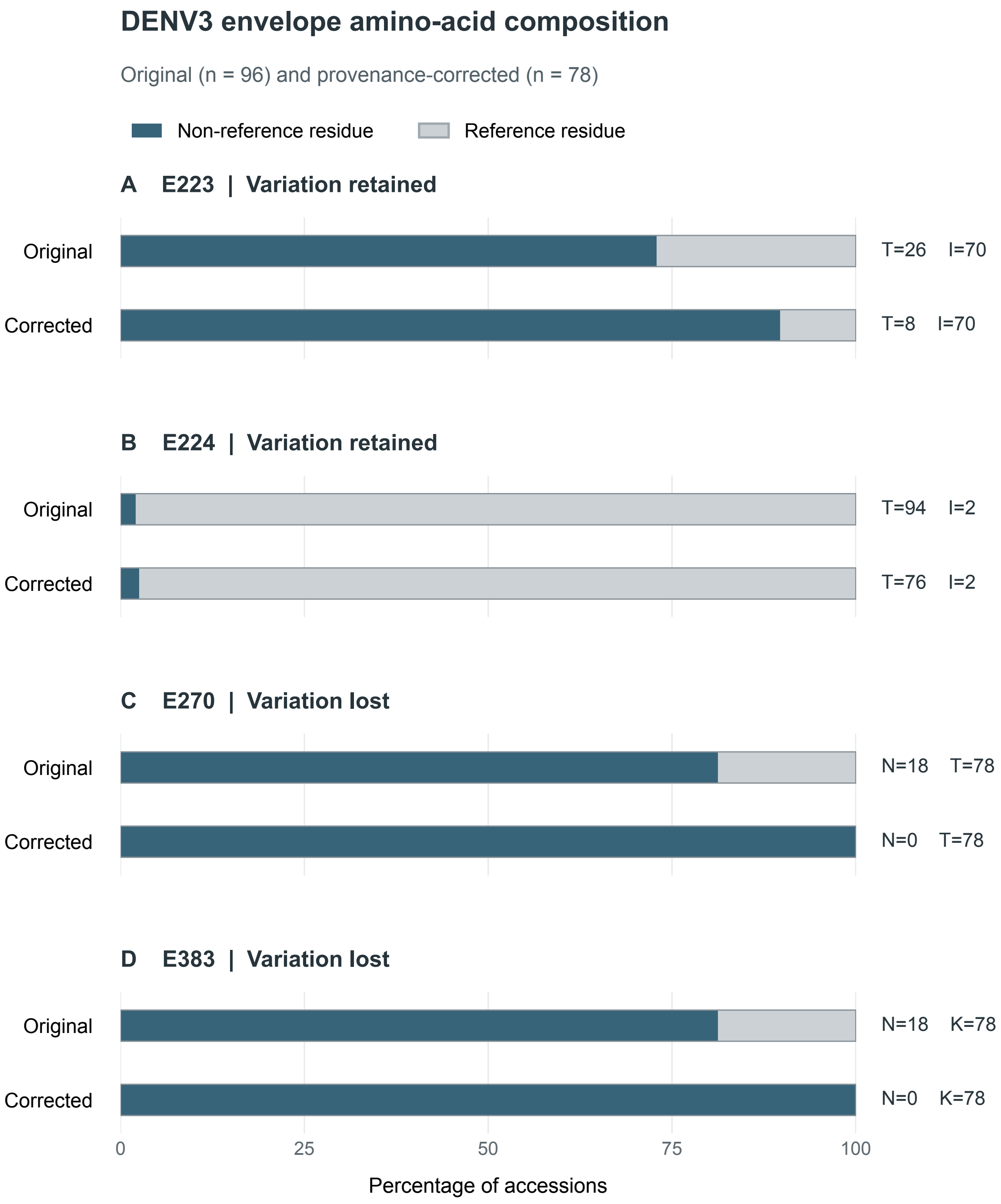

DENV3 envelope amino-acid composition
Original (n = 96) and provenance-corrected (n = 78)
Non-reference residue
Reference residue
A E223 | Variation retained
Original
Corrected
T=26 I=70
T=8 I=70
B E224 | Variation retained
Original
Corrected
T=94 I=2
T=76 I=2
C E270 | Variation lost
Original
Corrected
N=18 T=78
N=0 T=78
D E383 | Variation lost
Original
Corrected
N=18 K=78
N=0 K=78
0
25
50
75
100
Percentage of accessions
