## Supplementary Figure S2 - DENV3 variable-site classification matrix for "Provenance-aware handling of identical dengue genome records alters inferred envelope-protein variation in Bangladesh DENV-3"

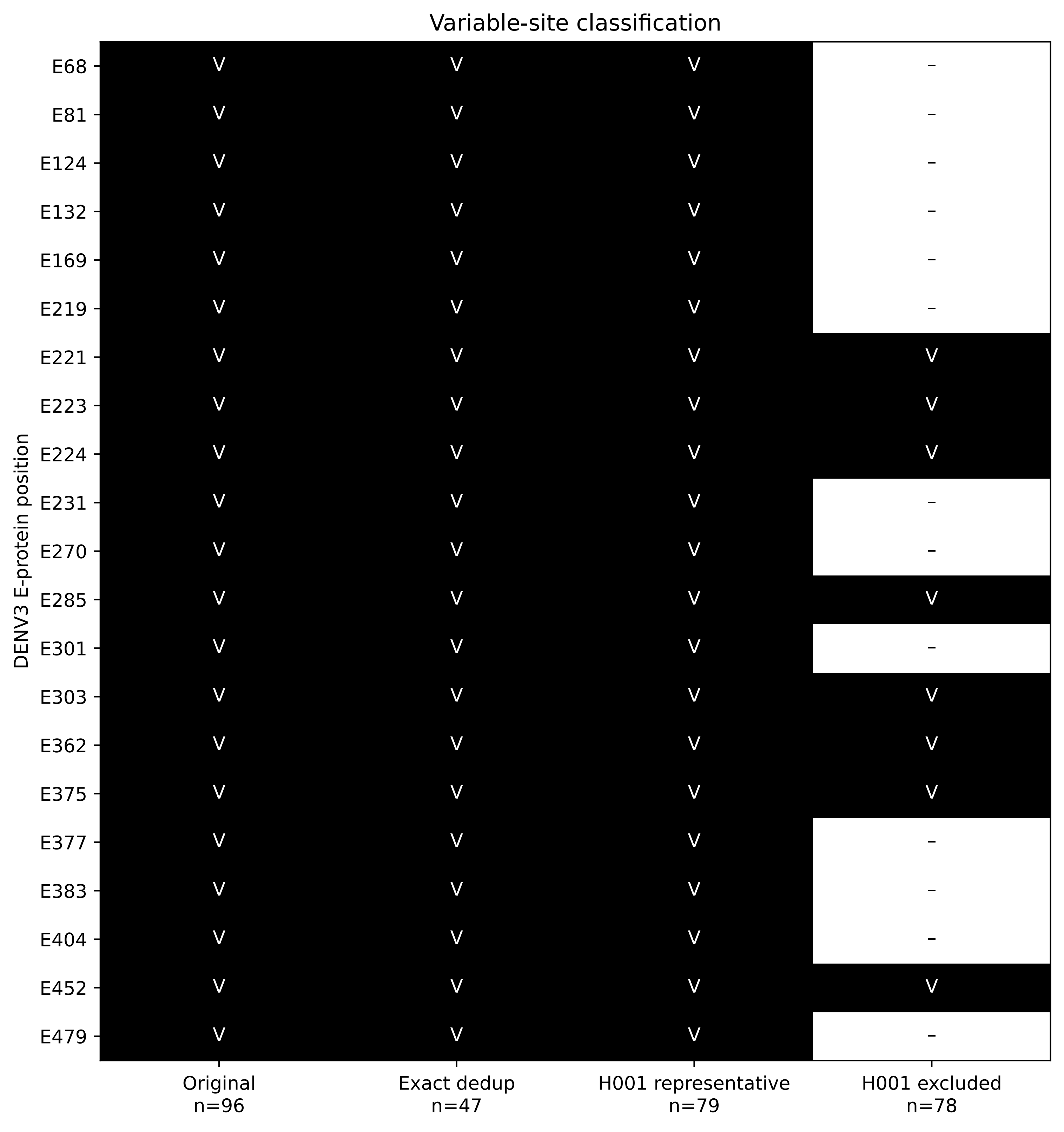
Variable-site classification
DENV3 E-protein position
E68
E81
E124
E132
E169
E219
E221
E223
E224
E231
E270
E285
E301
E303
E362
E375
E377
E383
E404
E452
E479
Original
n=96
Exact dedup
n=47
H001 representative
n=79
H001 excluded
n=78
